# DeepGSEA: Explainable Deep Gene Set Enrichment Analysis for Single-cell Transcriptomic Data

**DOI:** 10.1101/2023.11.03.565235

**Authors:** Guangzhi Xiong, Nathan John LeRoy, Stefan Bekiranov, Aidong Zhang

**Affiliations:** Department of Computer Science, University of Virginia, Charlottesville, VA, USA; Department of Biomedical Engineering, University of Virginia, Charlottesville, VA, USA; Department of Biochemistry and Molecular Genetics, University of Virginia, Charlottesville, VA, USA

**Keywords:** Gene Set Enrichment Analysis, Single-cell RNA sequencing, Explainable Artificial Intelligence, Phenotype Classification

## Abstract

Gene set enrichment (GSE) analysis allows for an interpretation of gene expression through pre-defined gene set databases and is a critical step in understanding different phenotypes. With the rapid development of single-cell RNA sequencing (scRNA-seq) technology, GSE analysis can be performed on fine-grained gene expression data to gain a nuanced understanding of phenotypes of interest. However, due to the extreme heterogeneity of single-cell gene expression, current statistical GSE analysis methods sometimes fail to identify enriched gene sets. Meanwhile, deep learning has gained traction in specific applications like clustering and trajectory inference in single-cell studies due to its prowess in capturing complex data patterns. However, its use in GSE analysis remains limited, primarily due to interpretability challenges. In this paper, we present DeepGSEA, an explainable deep gene set enrichment analysis approach which leverages the expressiveness of interpretable, prototype-based neural networks to provide an in-depth analysis of GSE. DeepGSEA learns the ability to capture GSE information through our designed classification tasks, and significance tests can be performed on each gene set, enabling the identification of enriched sets. The underlying distribution of a gene set learned by DeepGSEA can be explicitly visualized using the encoded cell and cellular prototype embeddings. We demonstrate the expressiveness of DeepGSEA over commonly used GSE analysis methods by examining their sensitivity and specificity with four simulation studies. In addition, we test our model on three real scRNA-seq datasets and illustrate the interpretability of DeepGSEA by showing how its results can be explained. The source code of DeepGSEA is available at https://github.com/Teddy-XiongGZ/DeepGSEA.

## 1 Introduction

Since its proposal, single-cell RNA sequencing (scRNA-seq) has excited researchers for its ability to capture gene signatures within the fundamental unit of life: single cells. By measuring the expression profiles of individual cells, researchers identified substantial cellular heterogeneity in gene expression that was not found in bulk RNA-seq data [28]. To deal with cellular heterogeneity in single-cell analysis, a number of deep learning (DL) methods have been proposed to model the complex gene expression distributions and in turn, handle canonical processing of scRNA-seq data, such as gene expression normalization [6], batch correction [27], clustering [13, 41], cell type identification [7, 18], and patient classification [15, 52].

An important downstream task for scRNA-seq data analysis is gene set enrichment (GSE) analysis. Gene sets are groups of genes characterized by common biological function, chromosomal location, or regulation [49]. GSE analysis identifies the enriched gene sets that are over-represented or under-represented in a large set of genes by comparing the gene expression in cells under different conditions, helping us to understand the mechanism underlying phenotypes or experimental conditions of interest [49]. For simplicity, in the remaining parts of this paper, we will use the term “phenotype” to refer to any of the following: different cellular phenotypes (e.g., tumor cells), experimental conditions (e.g., virus-infected cells), or phenotypes of patients (e.g., diseases) from which cells are collected that researchers want to compare. For different phenotypes, the associated genes in an enriched gene set may be over-represented or under-represented. Most existing GSE analysis methods are based on differentially expressed (DE) genes, either by selecting a pre-defined DE gene list or by calculating a DE score for each gene [14, 24, 33, 37, 43, 48]. While most of such methods were initially designed for GSE analysis on bulk RNA-seq data, many variants have been proposed in recent years to better accommodate scRNA-seq datasets. However, these DE gene-based approaches may under-utilize the complex distribution of gene expression profiles that a gene set could have [5]. This is because a distribution of gene signatures is reduced to a single score or a Boolean value about whether it appears in the DE gene list or not in such methods.

Another type of GSE analysis method is the multiple functional class scoring (FCS) method [34]. Multiple FCS methods treat all genes of a gene set as multivariate features and compare the distribution of expression levels across cells in high-dimensional space. Such approaches use statistical analyses to test if cells of different phenotypes can be distinguished in the high-dimensional space [5, 9, 12, 23]. While these methods have the potential to capture information about multivariate gene expression distribution exhibited by each gene set, their use of traditional statistical analysis methods limits these approaches from elucidating the complex transcriptional patterns that cells of different phenotypes may possess. To the best of our knowledge, at present there have been no attempts to use DL for multivariate GSE analysis, which may be attributed to the lack of interpretability of deep neural networks (DNNs). Though a number of post-hoc approaches have been proposed to explain a well-trained model [32, 44, 50], the opacity of DNNs still prevents end users from understanding how predictions are made from a given set of inputs.

With the rapid development of explainable artificial intelligence, a number of ante-hoc explainable methods have been proposed to build DL models that are both powerful and interpretable [17, 19, 40], among which prototype-based architecture is a promising branch that can provide case-based reasoning for trained models [7, 26, 52]. In this paper, we provide a solution to leverage the expressiveness of DNNs for GSE analysis while preserving interpretability for the prediction. Specifically, we propose DeepGSEA, a DL-enhanced GSE analysis framework that predicts the phenotype while summarizing and enabling visualization of complex gene expression distributions of a gene set by utilizing intrinsically explainable prototype-based DNNs, where each prototype of cells corresponds to a specific phenotype class of interest. Significance tests can be performed on each gene set, allowing for the screening of gene set enrichment. Moreover, the underlying distribution of a gene set learned by the neural network can be visualized using the learned cell embeddings and prototypes. With the design of a shared backbone encoder, DeepGSEA is able to learn the common encoding knowledge shared across gene sets, which is shown to improve the model’s ability to mine phenotype knowledge from each gene set by our ablation studies (Appendix C.2). We demonstrate the expressiveness of DeepGSEA in performing GSE analysis by examining its sensitivity and specificity in four simulation studies with comparisons to commonly used existing tools. We also test the performance of DeepGSEA in real applications with three different real-world datasets, on which we demonstrate DeepGSEA’s interpretability by visualizing the learned complex latent distributions and the gene expression of cells represented by each prototype. Our contributions are summarized as follows:

- To the best of our knowledge, DeepGSEA is the first DL-based gene set enrichment analysis approach that is able to learn the complex multivariate distribution of genes associated with an enriched gene set.
- DeepGSEA preserves interpretability by showing its underlying decision-making mechanism through visualizations of learned prototypes and cell embeddings.
- The interpretation given by DeepGSEA reveals how gene sets are enriched, which is beneficial in understanding data with multiple phenotypes or multiple cell subpopulations.

## 2 Related work

The majority of GSE analysis methods are based on DE genes, including over-representation analysis (ORA) and univariate functional class scoring (FCS) methods [34]. ORA determines the enrichment of a gene set by testing if the occurrence of its associated genes in a pre-defined list of DE genes is by chance [24, 43, 48], where the DE gene list can be determined by performing DE analysis on each gene. However, this kind of methods only utilize genes that are considered differentially expressed, and information on the remaining ones is ignored, which may result in a high false negative rate if the variation of gene expression across different phenotypes is subtle. Univariate FCS methods take all genes into consideration by assigning each gene a DE score for further analysis. In such approaches, the enrichment of a gene set is typically determined by combining DE scores of all genes [14, 33, 37]. But as is mentioned in [5], the information in the multivariate distribution of a gene set is significantly under-utilized by these DE gene-based GSE analysis approaches.

In addition to the above methods which are based on DE genes, there is a non-DE-gene-based category, which mainly contains multivariate FCS methods [34]. Multivariate FCS methods directly calculate an enrichment score for each gene set based on the expression data by considering the expression profiles of the associated genes as a multivariate feature. The recent development of such GSE analysis methods included Vision [9], which calculates a score for each cell by summarizing the expression of associated genes in the gene set, and SCPA [5], which uses a nonparametric graph-based statistical framework to compare multivariate distributions in high-dimensional data. The use of traditional statistical frameworks makes them less expressive in modeling complex heterogeneous distributions than neural networks.

In the domain of DL, the lack of interpretability in vanilla DNNs has drawn significant attention in recent years [17, 19, 40]. As the learning of prototypes can summarize the information of subpopulations in data and provide us with case-based reasoning for the downstream tasks, several recent groups have tried to incorporate prototypes in DNNs for interpretable decision-making. Li et al. [26] proposed to learn prototypes for image data in the latent space and make predictions of the image’s class using the distances from the image to different prototypes. They interpreted the model by looking at its learned weights on different prototypes for the classification task. For scRNA-seq data, we proposed ProtoCell4P [52] which leverages the interpretability of prototype-based models to perform intelligible patient classification. The model learns cell type-informed prototypes of cells and classifies each patient by summarizing prototype information within each cell. Based on our previous modeling of scRNA-seq data with prototypes, in this paper, we further use prototype-based DNNs to understand different gene set information in a cell and leverage their expressiveness and interpretability for GSE analysis.

## 3 Methods

### 3.1 Problem Formulation

GSE analysis is usually performed by traditional statistical tests. To that end, to incorporate deep learning into GSE analysis, we need to formulate it as a task that DNNs can handle. [12] pointed out the close connection between finding DE genes and predicting the clinical outcome, based on which we present

**Assumption 1** *Let* {𝒟 _1_, ⋯, 𝒟_*C*_} *be the gene expression distributions across cells of C different phenotypes in high-dimensional gene expression space and* {𝔇_1_, ⋯, 𝔇_*C*_} *be the distributions in the latent space after some non-linear mapping. A gene set is considered as enriched for C phenotypes if and only if we can find a mapping function that maps* { 𝒟_1_, ⋯, 𝒟_*C*_} *to* {𝔇_1_, ⋯, 𝔇_*C*_} *in the latent space such that a decision boundary can be found to distinguish cells according to the latent distributions*.

In other words, the significance of the enrichment of a gene set is closely related to how well a model can classify cells of different phenotypes by learning appropriate mapping functions and decision boundaries, which converts the GSE analysis to a classification problem. Based on the assumption, we formulate the training of DeepGSEA as a multi-task learning process, where each task is to perform an independent phenotype classification given the information of one gene set only.

Suppose the input scRNA-seq data contains the expression profiles of *N* genes for *n* cells of *C* different phenotypes, and our target is to analyze the enrichment of *T* gene sets in cells across different phenotypes. Given profiles of the *n* cells {**x**_1_, ⋯, **x**_*n*_} (**x**_*i*_ ∈ ℝ^*N*^, ∀*i* ∈ {1, ⋯, *n*}) and corresponding phenotype labels {*y*_1_, ⋯, *y*_*n*_} (*y*_*i*_ ∈ {1, ⋯, *C*}, ∀*i* ∈ {1, ⋯, *n*}), the model is encouraged to accurately predict the phenotype of each cell given the information from each gene set individually. For each input cell **x**_**i**_, the model should provide *T* similarity estimations 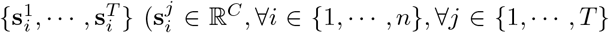 corresponding to the *T* gene sets of interest, each of which measures the similarity of the given cell to different phenotypes. Significance tests will be performed on each gene set by comparing the similarities for cells of different phenotypes, which returns a p-value indicating the significance of the enrichment of the gene set. Moreover, DeepGSEA can learn a set of *T* weights {*ω*_1_, ⋯, *ω*_*T*_} with a bias *ω*_0_ to linearly aggregate the results from different gene sets for more accurate predictions of cell phenotypes. The overall performance of phenotype classification reflects, in part, the appropriateness of the selected gene sets for understanding the phenotypes.

For the rest of this section, we use notations without the sample-subscript *i* ∈ { 1, ⋯, *n* }, in Section 3.2 to focus on the description of how the gene expression of one cell is processed. In Sections 3.3 and 3.4, the subscript *i* will be restored as all samples are involved in the model training and the GSE significance test.

### 3.2 Model Architecture

We present the overview of DeepGSEA in Fig. 1, which shows how the model performs gene set specific similarity estimation ({**s**^1^, ⋯, **s**^*T*^}) with the introduction of learnable prototypes and provides the final prediction **ŷ** of the phenotype by linearly aggregating the gene set information.

**Fig. 1:**
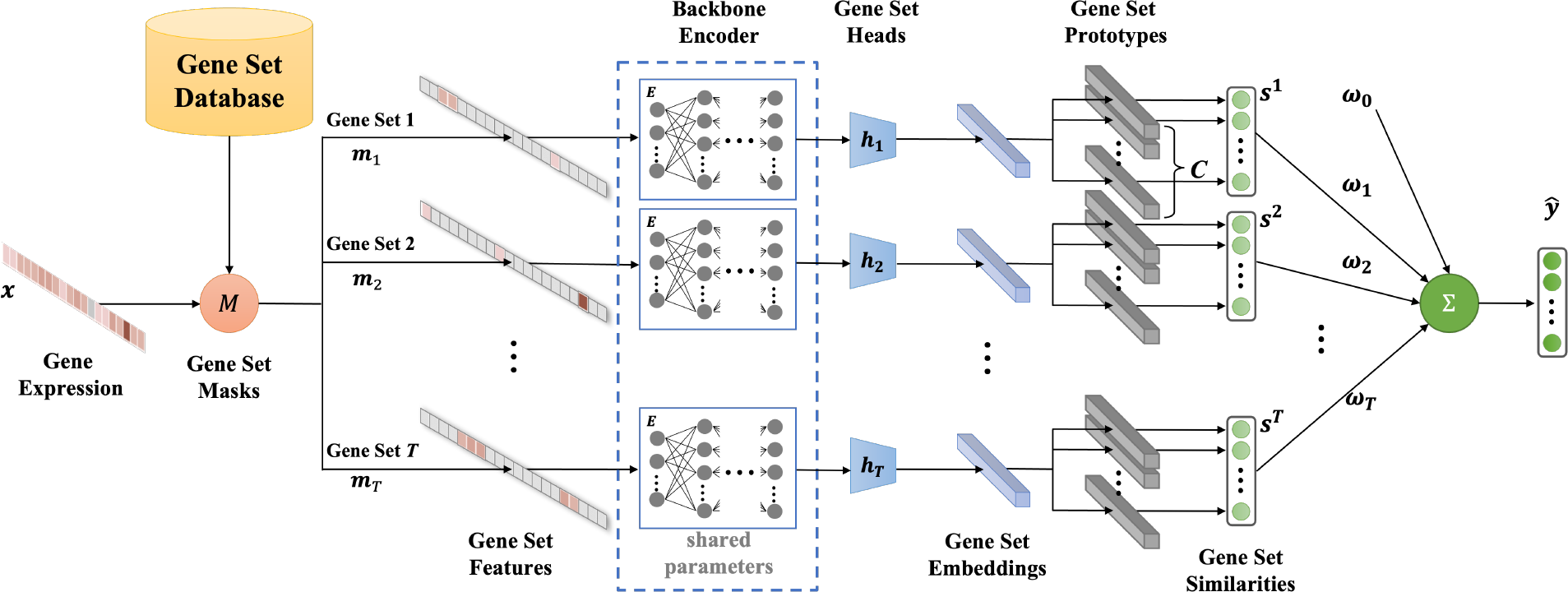
Overview of DeepGSEA.

#### Gene Set Feature Construction

In order to perform an independent and accurate analysis of each gene set, we first need to disentangle the information from different gene sets in the input cell. Given the annotations of associated genes for the *T* gene sets, we can construct a list of gene set masks {**m**_1_,⋯, **m**_*T*_} (**m**_*j*_ ∈ R^*N*^,∀*j* {1, ⋯, *T*}) such that the corresponding mask **m**_*j*_ of the *j*-th gene set has the value of 1 only at positions of the associated genes and 0 elsewhere. Given a cell *x*, the expression values of all unassociated genes of the *j*-th gene set can then be filtered out by applying

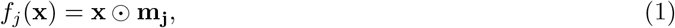

where ⊙ is the Hadamard product, and *f*_*j*_(**x**) is called the *j*-th gene set feature of the input. For a given cell **x**, we can disentangle the information of different gene sets and encode them as the gene set features {*f*_1_(**x**), ⋯, *f*_*T*_ (**x**)} for further analysis.

#### Backbone-head Gene Set Encoding

Since the real application of GSE analysis often involves the test of hundreds or thousands of gene sets, it is inefficient and impractical to build an independent complex neural network for each gene set. To tackle such a problem of inefficiency, we propose to use a “backbone-head” structure to share the knowledge among gene sets and make the model efficient. The logic behind such a design is, that the gene sets may overlap with each other, and therefore the knowledge of common genes could be shared by different gene sets. It is also shown in our ablation studies that the model’s learning of multiple gene sets really helps it to capture the phenotype information within each individual gene set.

Specifically, DeepGSEA is designed to learn a backbone encoder *E* and a list of gene set heads {*h*_1_,⋯, *h*_*T*_}, where *E* is a neural network that encodes the gene profile information and maps each input from the original gene expression space in ℝ^*N*^to the hidden space in ℝ^h_dim^ (h_dim is the dimension of the hidden space), and the gene set heads are one-layer fully connected models that map the vectors in the hidden space to cell embeddings in different gene set-specific latent spaces in ℝ^z_dim^ (z_dim is the dimension of the latent spaces), which output

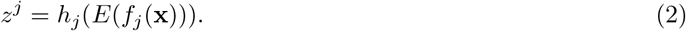

In this paper, *z*^*j*^ is called the *j*-th gene set embedding of the input, which should contain the well-encoded information of the cell’s gene profiles that are related to the *j*-th gene set. Since the heads for different gene sets are distinct, it results in diverse latent spaces for each of them, which preserves the heterogeneity of different gene sets in the encoding.

#### Prototype-based Similarity Measurement

As the latent space learned by a neural network can be extremely complex and obscure, its decision boundary for a classification task can be complicated and thus it is difficult to interpret what has been learned in the latent space. While preserving the expressiveness of DNNs in mapping the gene expression data to the latent embeddings, we propose to regularize the latent space by simplifying the form of decision boundaries and aligning it with prior biological knowledge for better interpretability. For each gene set, we have the following assumption:

#### Assumption 2

*On each gene set, for the cell distributions of C phenotypes* {𝒟_1_, ⋯, 𝒟_*C*_}, *we can find an appropriate mapping function that maps the original distributions in the gene expression space to* {𝔇_1_, ⋯, 𝔇_*C*_} *in the latent space, where each* 𝔇_*k*_ *(k* ∈ {1, ⋯, *C*}*) is a mixture of Gaussian distributions*.

This assumption follows the success of single-cell clustering studies where the assumption is widely taken [30, 31, 53]. On the basis of this assumption, we propose to learn additional global latent vectors as the centers of different Gaussian distributions in the latent space. Formally, for the *j*-th gene set and its latent distribution of the *k*-th phenotype, DeepGSEA learns a list of latent vectors 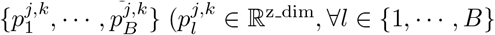 corresponding to the centers of *B* Gaussian distributions in the mixture model, which we call prototypes, as each of them represents a cell subpopulation in the latent space of gene sets. With the introduced prototypes, we simplify the decision boundary for the phenotype classification task by letting the model predict a cell’s phenotype via assigning it the label of the most “similar” prototype. Given the *j*-th gene set embedding of a cell *z*^*j*^, its similarity to the *k*-th phenotype is defined as

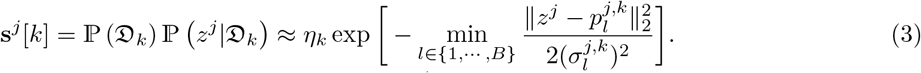

where **s**^*j*^[*k*] stands for the *k*-th entry in the similarity vector **s**^*j*^. *η*_*k*_ is the estimated proportion of cells of the *k*-th phenotype and 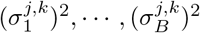 are learnable variances for corresponding prototypes. The probability of the given cell being classified into the *k*-th phenotype group can then be computed as

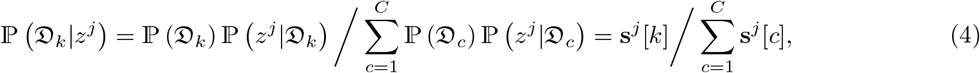

The final prediction of a cell’s phenotype can be provided by linearly combining the encoded information from each gene set as

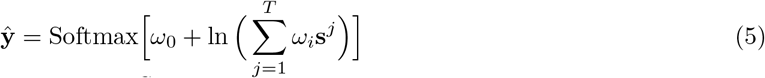

where 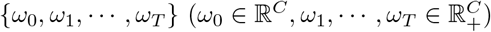 are the bias and weights to be learned.

### 3.3 Loss Computation and Model Training

The training objective of a DNN is crucial for its training as it will guide the model to learn correct knowledge as desired. In the training of DeepGSEA, a series of loss functions are carefully designed for the learning of prototypes and embeddings. First, we encourage the model to provide a good prediction of phenotypes with the estimated probabilities based on each gene set. Given *n* cells {**x**_1_,⋯, **x**_*n*_}, the model is trained to minimize the cross entropy losses

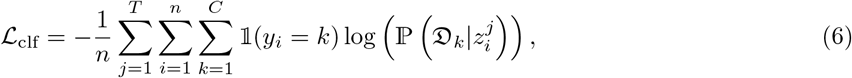

where 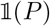 is 1 if *P* is true and 0 otherwise. While the minimization of ℒ_clf_ encourages the model to well predict phenotypes based on each gene set information, we also train it to minimize the cross entropy loss given by the final prediction

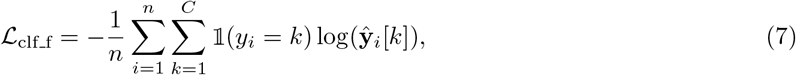

which will guide the model to learn proper *ω*_0_, *ω*_1_, ⋯, *ω*_*T*_.

ℒ_clf_ and ℒ_clf_f_ generally encourage DeepGSEA to differentiate cells of different phenotypes in the latent space. However, the hypothesis space of DNNs is so large that learned prototypes may work well in classification but are far from the cells they represent in the latent space, which can result in undefined behaviors when outliers are entered. Therefore, we should encourage the prototypes to be as close to the cells they represent and encourage large pairwise distances of prototypes so that different cell subpopulations can be represented. Formally, the loss for the pairwise prototype distance is defined as

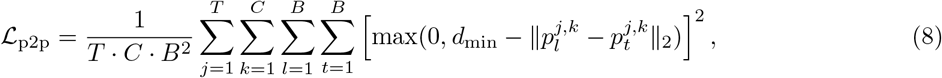

where *d*_min_ is the minimum acceptable distance between prototypes which is defined as 1 in our implementation. The long distances between cells and corresponding prototypes are penalized by the minimization of the cell-to-prototype loss, which is defined as

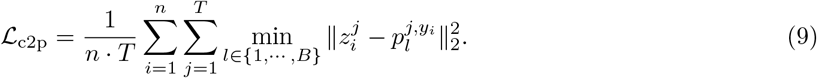

As can be observed, ℒ_c2p_ encourages the model to minimize the distance from each cell to the closest prototype with respect to the same phenotype. Since the learned prototype may be located at the border of the cell subpopulation it represents, in addition to ℒ_c2p_, we also encourage the model to minimize the distance from each prototype to the center of cells to which it is the closest, i.e., cells it represents

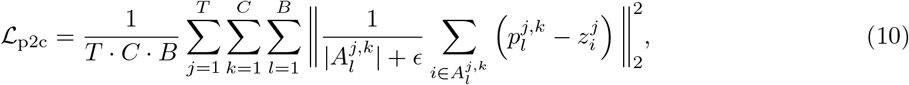

where 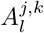 is defined as

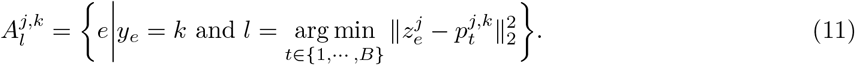

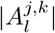 denotes the size of 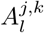 and *ϵ* is an extremely small value, which is set as 10^*−*16^ in our experiments.

As various cell types may be present in scRNA-seq data, the diversity of different cell subpopulations can lead to the heterogeneity of cells in a dataset. With the existing cell clustering and cell type annotation methods [13, 41], it is easy to obtain labels or pseudo-labels for each cell in the scRNA-seq data, which is beneficial for the understanding of cell heterogeneity. Given such labels, DeepGSEA can incorporate the additional biological knowledge into its training for better performance and interpretability by modifying the loss functions relating to prototype learning. Specifically, *B* in Formula 3 will be set as the number of biological labels with each of them corresponding to a particular label (e.g., a cell type). Instead of merely penalizing the pairwise prototype distance, we encourage DeepGSEA to well predict the biological label of the input cell given its distances to different prototypes. Suppose the corresponding additional labels of the cells are {*q*_1_, ⋯, *q*_*n*_} (*q*_*i*_ ∈ {1, ⋯, *B*}, ∀*i* ∈ {1, ⋯, *n*}). Formally, ℒ_p2p_ is re-defined as

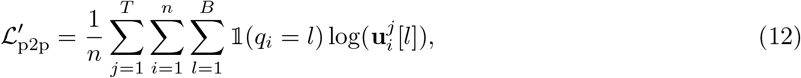

where 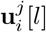 is probabilities of the *i*-th cell being classified into the *l*-th label given the information of the *j*-th gene set, which is formulated as

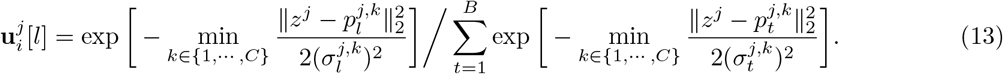

In the meantime, ℒ_c2p_ and ℒ_p2c_ can also be updated to focus on the distance between each prototype to cells it represents that have both the same phenotype and the same label. ℒ_c2p_ is now re-formulated as

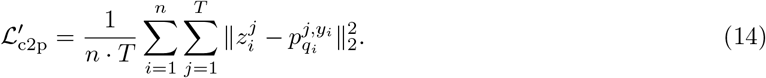

And ℒ_p2c_ can be re-formulated as

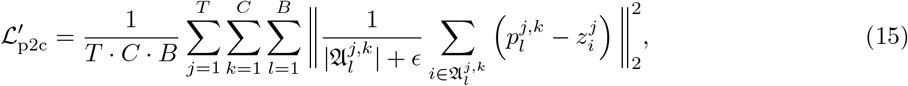

where 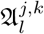 is defined as

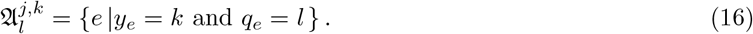

The overall training objective is minimizing a linear combination of all loss functions above. When DeepGSEA is trained without additional biological labels of cell subpopulations, the overall loss function is

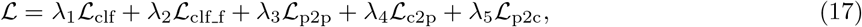

where {*λ*_1_, ⋯, *λ*_5_} are the weights for different losses. Given additional labels, the loss function can be

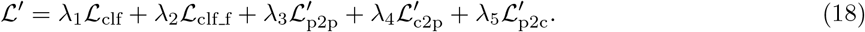

As the ℒ_clf_f_ is designed to learn appropriate *ω*_0_, *ω*_1_,, *ω*_*T*_ to aggregate results from different gene sets, which relies on the quality of the encoding of gene set information, in our implementation, a two-stage training paradigm is used, where *λ*_2_ is explicitly set as 0 in the first stage and is restored to the preset value in the second stage. This allows the model to focus on the learning of gene set encoding first and then tune the parameters for the final prediction based on the well-trained gene set encoders.

### 3.4 Significance Test of Gene Set Enrichment

We propose to use the Mann-Whitney U test [35] to test the significance of GSE given the results from the model. Let **MWU**(·, ·) be the function that takes two lists of values and outputs a p-value for the Mann-Whitney U test. For the significance test of *C* phenotypes, we perform the Mann-Whitney test on estimated cell similarities to each phenotype class and then combine the p-values using Fisher’s method [10] to obtain a final p-value for the *j*-th gene set. Specifically, for the *k*-th phenotype class, DeepGSEA can provide the similarity score of each cell to this class using Formula 3. Ranking the scores of all cells, we test whether the average ranking of cells with the *k*-th phenotype is equal to the average ranking of cells that do not have the *k*-th phenotype, with a p-value reflecting the probability that the score of a random cell with the *k*-th phenotype is smaller than the score of a random cell with other phenotypes, which can be formulated as

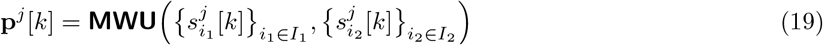

where *I*_1_ = {*e*|*y*_*e*_ = *k*} stands for the set of indices of cells with *k*-th phenotype and *I*_2_ = {*e*|*y*_*e*_ ≠ *k*} contains the indices of cells with other phenotypes. **p**^*j*^ is a list of p-values whose entries are computed based on the similarity measurement corresponding to each phenotype. Since they share the same null hypothesis that the mean ranks for scores in two groups are equal, we can use Fisher’s method to summarize the p-values and obtain one p-value for the GSE analysis of the *j*-th gene set:

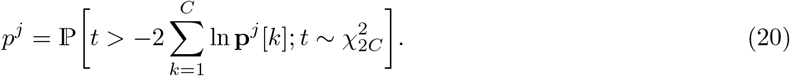

For the test of multiple gene sets, as is frequently required in real applications, we control the false discovery rate by applying the Benjamini-Hochberg correction [4] to the p-values. For the *K*-fold validation which is commonly used in the evaluation of DNNs, Pearson’s method [39] is used to combine the p-values given by models trained in different runs. Unlike Fisher’s method which is sensitive to the smallest p-value, Pearson’s method is more sensitive to the largest one when combining the p-values [16].

An overall algorithm of DeepGSEA for GSE analysis is provided in Appendix A.

## 4 Experiments

### 4.1 Datasets and Baselines

We first benchmark the performance of DeepGSEA against other commonly used approaches through four simulation studies. Moreover, three real-world datasets are selected to test the interpretability of DeepGSEA in real applications.

#### Baseline GSE Analysis Methods

The baseline GSE analysis methods we choose are those commonly used for single-cell transcriptomic data, including AUCell [1], GSVA [14], ssGSEA [3], Vision [9], SCPA [5], and Z scoring that are either based on DE genes or based on the multivariate distribution of each gene set for GSE analysis.

#### Simulated scRNA-seq Datasets

We follow [5] to simulate scRNA-seq datasets with 1000 cells of 2 different phenotypes using Splatter [54]. There are four designed experiments in our simulation, with variations on 1) the size of differential expression (log fold change) of genes in a selected gene set, 2) the probability that a gene will be differentially expressed in a selected gene set, 3) the number of cells in the dataset, and 4) the proportion of positive samples (e.g., the proportion of cells in the first group) in the dataset. P-values for the enrichment of the selected gene set are measured to test the sensitivity of different GSE analysis approaches. We also report the p-values for the enrichment of another irrelevant gene set that has no shared associated genes with the selected one to evaluate the specificity of DeepGSEA and other methods. The implementation details of the simulation can be found in Appendix B.1.

#### Real-world scRNA-seq Datasets

We select three real-world scRNA-seq datasets to evaluate the interpretability of DeepGSEA in real applications, including the analysis of glioblastoma [56], influenza [42], and Alzheimer’s disease [55]. The detailed descriptions of the datasets and how we pre-process them are provided in Appendix B.2. For each dataset, we determine the candidate gene sets to analyze by identifying gene sets that overlap with the list of genes in the given dataset. Specifically, gene sets with at least one associated gene present in the scRNA-seq dataset will be taken into account. Two different sources of gene sets are considered, which are Gene Ontology (GO) [2, 8] and Pathway, a constructed collection of Hallmark [29], KEGG [20], and Reactome [11]. In particular, to control the granularity of analyzed gene sets, we select only terms at the same level in GO for analysis. In our experiments, biological processes at level 5 are chosen for the GSE analysis study. A summary of the real-world datasets we use is provided in Table 1.

**Table 1:** Real-world datasets for the evaluation of DeepGSEA.

| Dataset Name | Num. of groups | Num. of cells | Num. of genes | Num. of cell types | Num. of GO terms | Num. of pathways |
| --- | --- | --- | --- | --- | --- | --- |
| glioblastoma [56] | 2 | 132 | 21209 | – | 1870 | 1887 |
| influenza [42] | 3 | 18640 | 25379 | – | 1870 | 1889 |
| alzheimer [55] | 2 | 16692 | 2766 | 13 | 1722 | 1718 |

### 4.2 Expresiveness of DeepGSEA on Simulated scRNA-seq Data

DeepGSEA and baseline methods are run on all simulated datasets and the results are summarized in Fig. 2 and 3. The x-axis stands for the value of parameters that vary in each simulation, and the y-axis shows the negative log-transformed p-values for GSE analysis provided by each method. The larger the value on the y-axis, the more significant the enrichment of the selected gene is found by a GSE analysis approach.

**Fig. 2:**
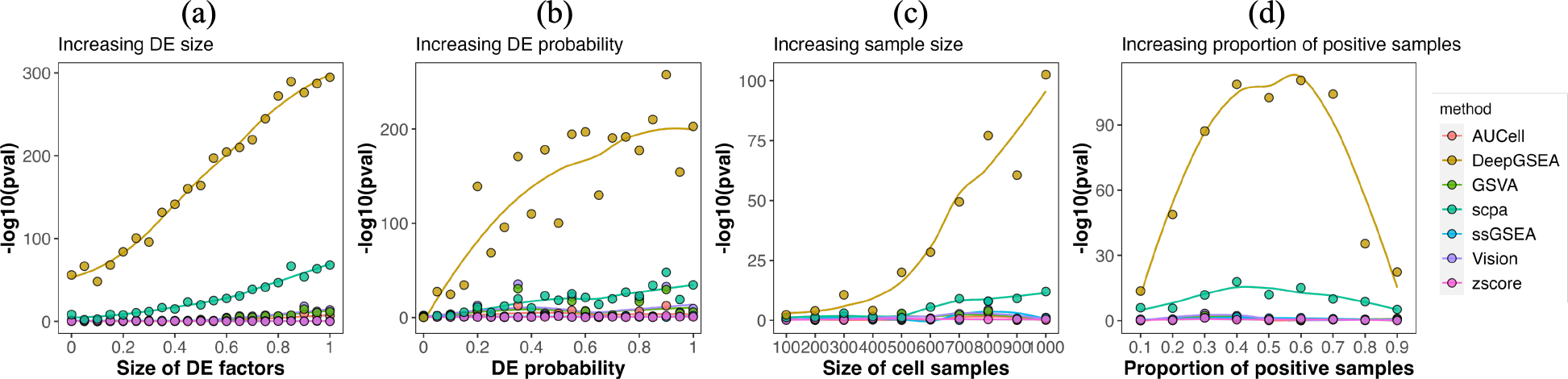
Sensitivity test: significance of the enrichment of the relevant gene set.

**Fig. 3:**
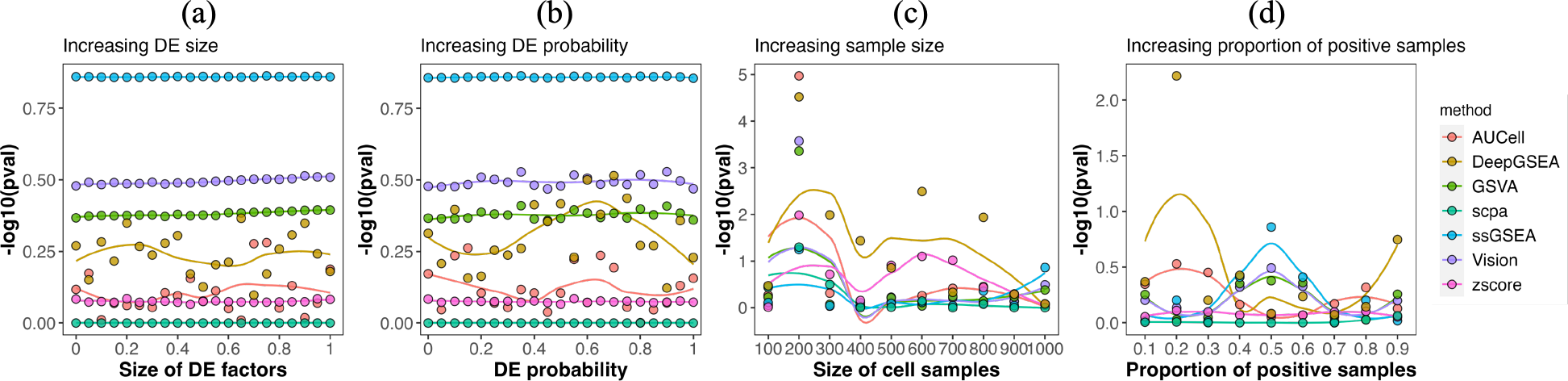
Specificity test: significance of the enrichment of the irrelevant gene set.

Fig. 2 displays the results of the sensitivity test, showing the significance of enrichment of the selected gene set estimated by different GSE analysis methods in the four simulation studies. It can be clearly observed that DeepGSEA is more sensitive to the GSE signals than all other approaches in comparison. In particular, Fig. 2 (a) and (b) show that DeepGSEA is able to precisely capture the difference between gene profiles of cells from different groups even when the variation is subtle. It should be noted that the DE size of 0 in Fig. 2 (a) corresponds to a non-trivial differential expression of the selected gene set, which is explained in detail in Appendix B.1. Thus, it is surprising but reasonable to see that DeepGSEA can provide a significant p-value (5.8×10^*−*57^) that is much smaller than other methods (e.g., 4.0 × 10^*−*9^ by SCPA, 0.05 by ssGSEA) even when the DE size is 0. As the signal of variation is amplified, our model will consider the gene set as enriched more positively and confidently than all other baseline approaches we compare [1, 3, 5, 9, 14]. Meanwhile, Fig. 2 (c) and (d) demonstrate that our method is able to identify potential enriched gene sets even in extreme cases where the data has very few or unbalanced samples. It is also revealed that DeepGSEA can output more significant results when dealing with abundant balanced data, which is becoming practical in real applications with the development of scRNA-seq technologies.

Fig. 3 shows the results of enrichment analysis of the irrelevant gene set by different GSE analysis methods. We can observe from the results that the specificity of DeepGSEA is comparable to other commonly used approaches in most cases, as GSE analysis of the irrelevant gene set always returns a high p-value. However, Fig. 3 (c) and (d) also suggest that our approach may be more sensitive in capturing the spurious correlation between phenotypes and gene expressions than other methods when the dataset is small or unbalanced, which can result in more false discoveries. But as mentioned above, the rapid development of scRNA-seq research has allowed us to utilize more single-cell transcriptomic data for the analysis of phenotypes, in which scenarios DeepGSEA can play a considerable role.

Results of DeepGSEA on simulated data show that our model is expressive enough to capture gene profile differences in enriched gene sets with better sensitivity than the compared baselines and comparable specificity, which demonstrates its usefulness in applications.

### 4.3 Interpretability of DeepGSEA on Real-world scRNA-seq Datasets

In addition to usefulness, DeepGSEA is trustworthy as it is intrinsically explainable with prototype-based designs. After performing GSE analysis on three real-world scRNA-seq datasets with DeepGSEA, we demonstrate its interpretability by providing explanations for enriched gene sets it identifies. Additional results about its phenotype classification performance and ablation studies are included in Appendix C.

#### Interpretation on Glioblastoma Data

Despite the small number of cells in the glioblastoma dataset, DeepGSEA can effectively identify enriched gene sets with small p-values. Fig. 4 presents the visualization of latent spaces of selected gene sets using UMAP [36], where each dot corresponds to the embedding of a cell and the stars represent the prototypes of cells corresponding to different phenotypes. The color of each dot or star reflects the biological semantics displayed in the legend. The GO term “acid secretion” is the biological process whose associated genes are found to be significantly enriched (*p* = 1.6× 10^*−*26^) in the glioblastoma dataset, which is consistent with the existing discovery that the waste product of glioblastoma cell fermentation will acidify the microenvironment [47]. It is shown by Fig. 4 (a) that cancer cells and neural cells can be clearly separated in the embedding space of “acid secretion”. Similar analysis can be performed on pathways that are found to be enriched. Fig. 4 (b) shows the gene expression in different groups tends to differ with respect to the pathway “vldlr internalisation and degradation” (*p* = 1.6 × 10^*−*34^), which corresponds to the discovery that VLDLR is upregulated in glioblastomas [45] and thus associated genes of “vldlr internalisation and degradation” should be under-represented in the cancer cells. We also present the visualization of another pathway (Fig. 4 (c)) that is not found to be enriched (*p* = 1.0). As can be seen, the latent space of this pathway looks chaotic, with a mixture of cells of different phenotypes.

**Fig. 4:**
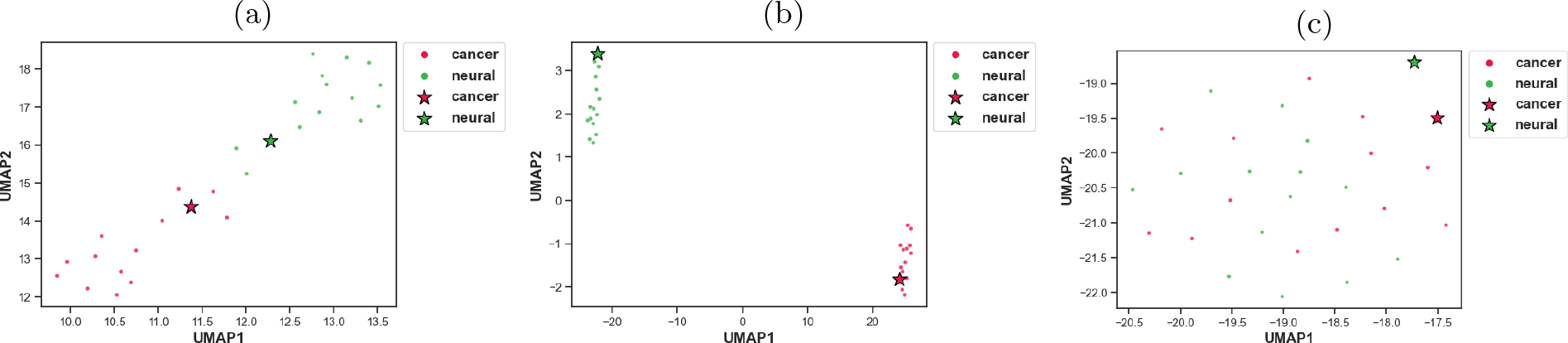
Visualization for latent spaces of different gene sets in the glioblastoma dataset: (a) “acid secretion” from GO, (b) “vldlr internalisation and degradation” from Pathway, (c) “relaxin receptors” from Pathway.

In addition to visualizing the latent space, we can also plot the heatmap for cells around each prototype to see what they really represent and gain a nuanced understanding of phenotype differences at the level of single genes. Fig. 5 shows the heatmap of cells around the learned prototypes in the latent space of “acid secretion”, where each column corresponds to an associated gene for the pathway and each row represents a cell that is close to either prototype. The color of each unit reflects the log-transformed expression counts of a gene (columns) in a given cell (row). We can observe from the figure that “P2RX7” is over-represented in cancer cells while “OXTR” is under-represented, corresponding to related biological discoveries [21, 25, 46].

**Fig. 5:**
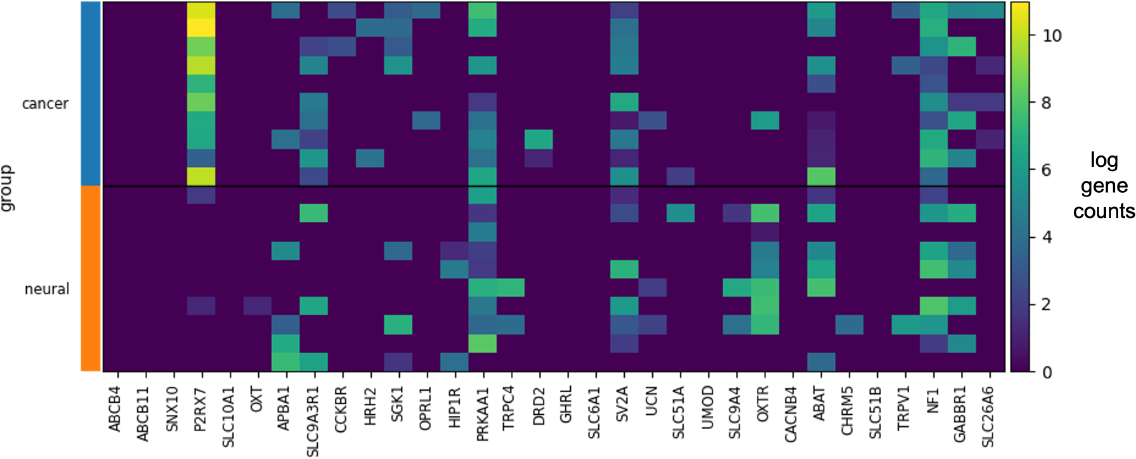
Heatmap of log gene counts of associated genes in the pathway (columns) for cells (rows) that are close to each prototype in the latent space of “acid secretion”, where blue corresponds to low expression and yellow corresponds to high expression.

#### Interpretation on Influenza Data

DeepGSEA’s ability to interpret gene sets with visualization can be beneficial in the analysis of data with multiple phenotypes, as it is able to show how a gene set is enriched across different groups. Fig. 6 (a) and (b) present visualized high-dimensional spaces of “translation”, with the gene set features and gene set embeddings of cells (in Fig. 1) as the data points, respectively. While cells are mixed in the original high-dimensional space of pre-processed gene expression, they can be well separated in the latent space by utilizing the gene set information. Though three clusters are clearly identified in Fig. (b), we can observe that the mock group and the MOI0.2 group are close to each other with some intersections while MOI2.0 is distant from both of them in the latent space, indicating that “translation” pathway genes in cells infected by the influenza virus at MOI of 2.0 are more differentially expressed compared to the mock ones than those in cells infected by the virus at MOI of 0.2. The visualization of another enriched gene set is shown in Fig. 6 (c) . Unlike “translation” on which cells of the three groups can be distinguished, the pathway “ho gtpases activate cit” only separates the mock group from the other two in the latent space, while cells in the two infected groups are mixed and cannot be separated using this gene set.

**Fig. 6:**
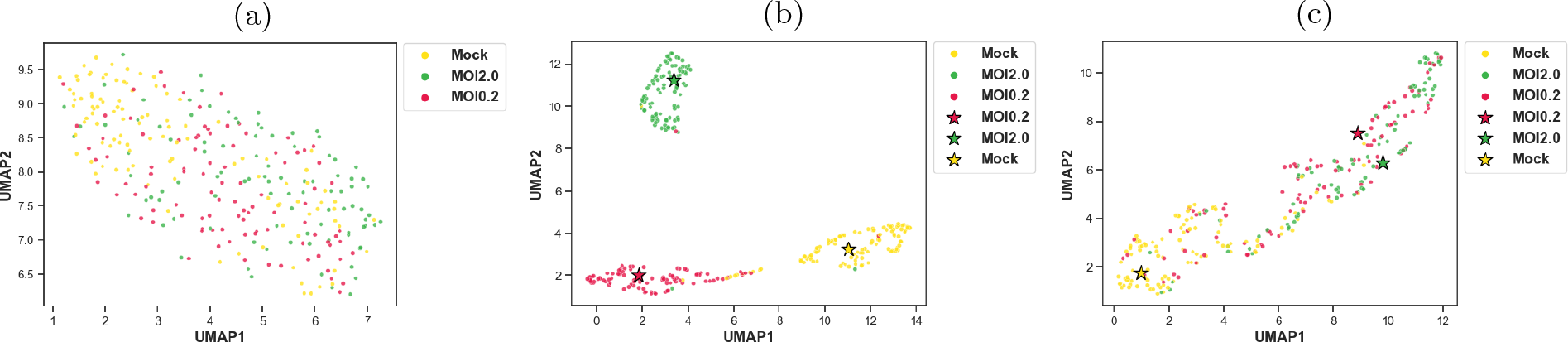
Visualization of cells in the influenza dataset: (a) “translation” information before encoding, (b) “translation” information after encoding (b) “rho gtpases activate cit” information after encoding.

Additionally, the phenotype similarities of each cell can be visualized on the same figure with colors indicating the estimated similarities. Fig. 7 shows similarities of all cells to each phenotype. We can see that cells around each prototype are correctly recognized by the model with high similarity to the corresponding phenotype, whereas cells located at the boundary of “Mock” and “MOI0.2” have less similarity to both groups, as they are less typical and intermixed. The visualization of DeepGSEA’s similarity estimation provides us with an intuitive understanding of what underlying distributions are learned by the model.

**Fig. 7:**
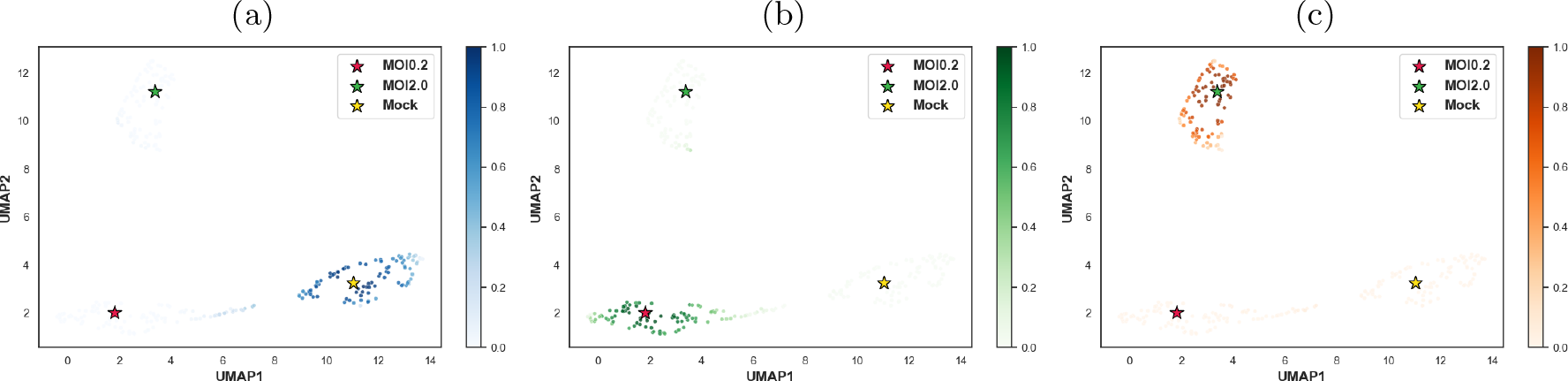
Visualization for learned similarities of cells in the influenza dataset based on “translation”: (a) similarity to “Mock”, (b) similarity to “MOI0.2”, (c) similarity to “MOI2.0”.

#### Interpretation on Alzheimer’s Disease Data

For the alzheimer dataset, DeepGSEA is trained with the cell type annotations indicating different cell subpopulations in the data. Fig. 8 (a) and (b) show the visualized latent space of the biological process “generation of neurons” with dots marked by cell types and phenotypes, respectively. It can be observed that DeepGSEA can effectively learn the knowledge of phenotypes in the gene set while preserving the information of cell types. In particular, the information of each phenotype is captured by learning phenotype-specific prototypes within each cell type, which returns a learned Gaussian mixture model for the associated distribution. Fig 8 (c) shows the prediction of cells given by DeepGSEA, with the color of dots in the plot reflecting the model’s predicted probability of the disease group for each cell. We can see that the complex distribution of phenotypes is accurately learned by DeepGSEA with the help of the additional cell type knowledge. For example, DG cells in the disease group are surrounded by cells in the control group. By learning a prototype to represent such a subpopulation, these DG disease cells can be correctly identified without being overwhelmed by cells nearby.

**Fig. 8:**
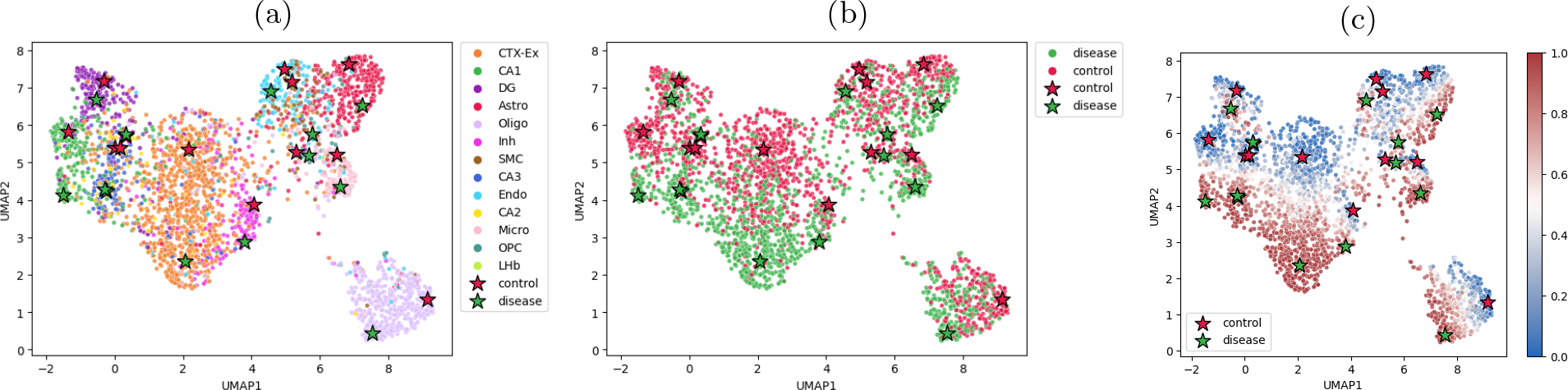
Visualization for the latent space of “generation of neurons” in the alzheimer dataset: (a) cells marked by cell types, (b) cells marked by phenotypes, (c) cells marked by predicted probabilities of “disease”.

Moreover, DeepGSEA can provide a nuanced interpretation of the learned distribution for each cell type. The distributions of two cell types in the latent space of “generation of neurons” are visualized in Fig. 9. The CA1 cells of different phenotypes can be clearly distinguished by DeepGSEA, corresponding to the biological discovery that hippocampal CA1 neurons are impaired in Alzheimer’s disease [22, 38, 51]. However, the OPC cells of different phenotypes are rather mixed in the latent space, suggesting that they are less distinguishable with the current gene set. Such a nuanced interpretation can help users understand the learned distribution within each gene set more easily, especially when the overall distribution is complex.

**Fig. 9:**
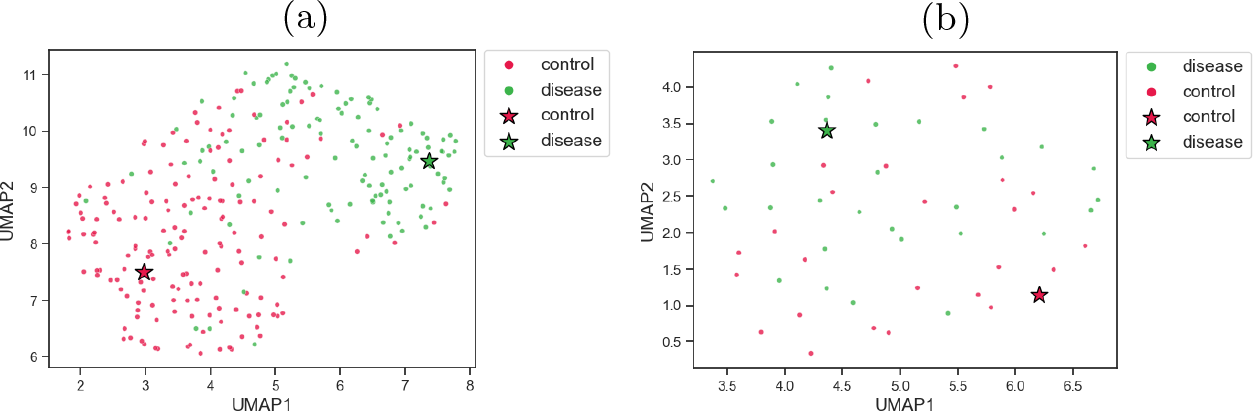
Visualized subspaces of “generation of neurons” for different cell types: (a) CA1 cells, (b) OPC cells.

## 5 Conclusion

We presented an explainable deep gene set enrichment analysis (DeepGSEA) method for in-depth GSE analysis on single-cell transcriptomic data, which utilizes interpretable prototype-based neural networks to model complex distributions that gene sets can exhibit in real data. Compared to existing commonly used approaches, DeepGSEA is much more sensitive while preserving comparable specificity, indicating DeepGSEA is highly expressive in capturing gene profile differences across cells of various phenotypes. Experiments on real-world datasets demonstrate that DeepGSEA is also interpretable, as one can always explain how a gene set is enriched by visualizing the latent distributions and gene set projected expression profiles of cells around the learned prototypes. A high degree of expressiveness and interpretability enhance the usefulness and trustworthiness of DeepGSEA, making it a powerful tool for real-world GSE analysis.

## A Algorithm of DeepGSEA

Algorithm 1 presents the pseudocode of using DeepGSEA for GSE analysis. A 5-fold validation is implemented for all experiments in our study.

### Algorithm 1

The algorithm of DeepGSEA for the analysis of gene set enrichment

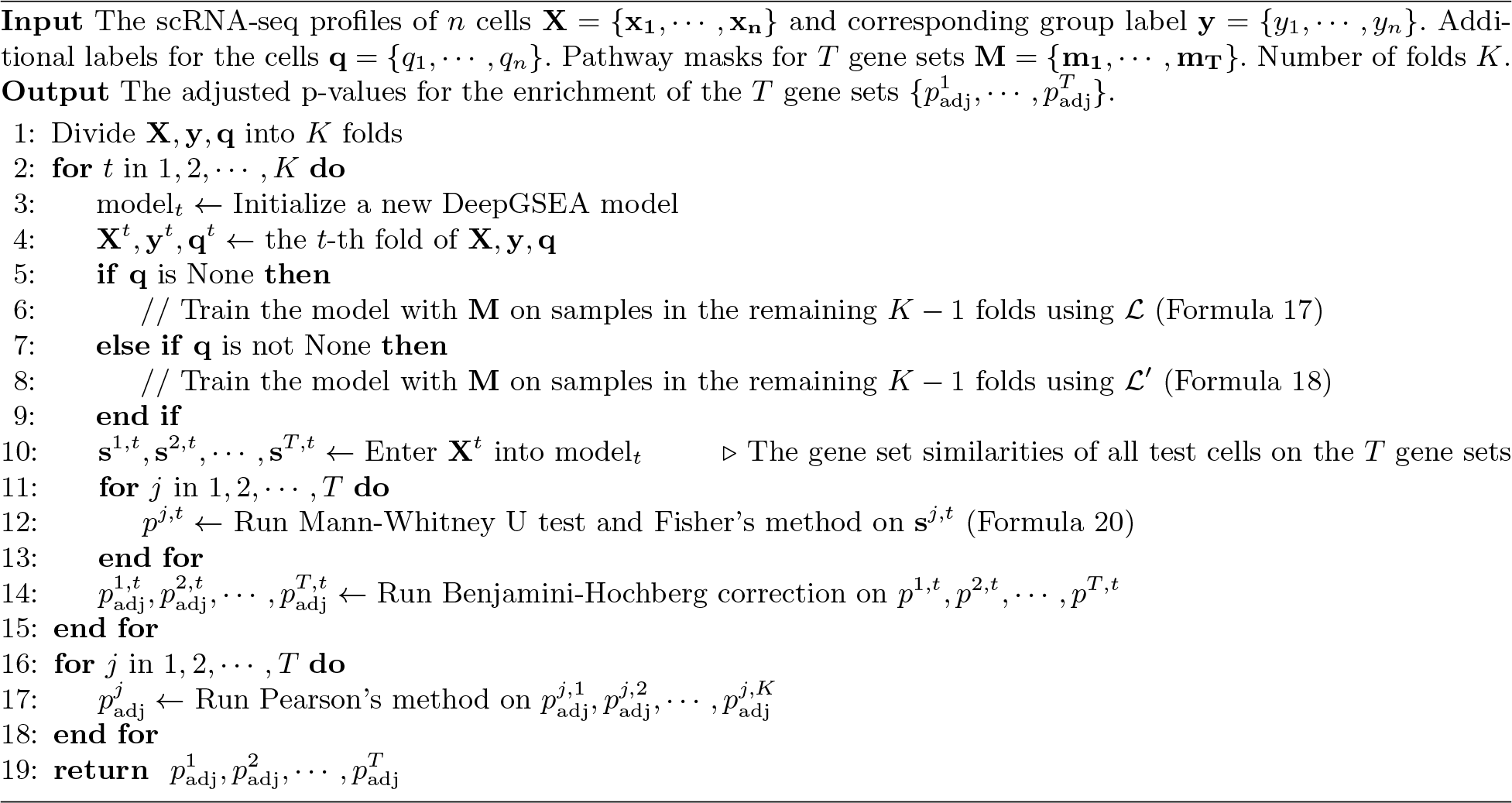

## B Implementation Details

### B.1 Design of Simulated Datasets

“HALLMARK COMPLEMENT” is selected as the enriched gene set for the sensitivity test in our simulation study, and “HALLMARK NOTCH SIGNALING” is chosen as the irrelevant gene set for the specificity test. Following [5], we first use Splatter to estimate simulation parameters from their real scRNA-seq data^1^. The definitions of some key parameters are shown as follows^2^

- batchCell: *the number of cells in each batch*. In our simulations, it corresponds to the number of all cell samples in a simulated scRNA-seq dataset.
- group.prob: *a vector giving the probability that cells will be assigned to groups*. In our study, group.prob is a vector with non-negative entries whose sum is 1.
- de.prob: *this parameter controls the probability that a gene will be selected to be differentially expressed*.
- de.facLoc: *Differential expression factors are produced from a log-normal distribution. Changing these parameters can result in more or less extreme differences between groups*. When de.facLoc is 0, the DE factors for each gene will not be strict 1, since the default value of the parameter de.facScale for the log-normal distribution is 0.4 instead of 0. Thus, the genes can be either over-represented or underrepresented, which makes the differential expression non-trivial.

Then the default parameters for the background data and the selected gene set are updated according to Table 2.

**Table 2:** Default parameters for simulated background data.

| Background |  |  |  | HALLMARK_COMPLEMENT |  |  |  |
| --- | --- | --- | --- | --- | --- | --- | --- |
| batchCells | group.prob | de.prob | de.facLoc | batchCells | group.prob | de.prob | de.facLoc |
| 1000 | (0.5,0.5) | 0.2 | 0.2 | 1000 | (0.5,0.5) | 0.3 | 0.3 |

The simulation studies are performed by varying the different parameters to see if the model performance will change. The variations of parameters in the four experiments are listed as follows:

1. Variation on “de.facLoc” for “HALLMARK COMPLEMENT”: {0.00, 0.05, 0.10, 0.15, 0.20, 0.25, 0.30, 0.35, 0.40, 0.45, 0.50, 0.55, 0.60, 0.65, 0.70, 0.75, 0.80, 0.85, 0.90, 0.95, 1.00}.
2. Variation on “de.prob” for “HALLMARK COMPLEMENT”: {0.00, 0.05, 0.10, 0.15, 0.20, 0.25, 0.30, 0.35, 0.40, 0.45, 0.50, 0.55, 0.60, 0.65, 0.70, 0.75, 0.80, 0.85, 0.90, 0.95, 1.00}.
3. Variation on “batchCells” for both background and “HALLMARK COMPLEMENT”: {100, 200, 300, 400, 500, 600, 700, 800, 900, 1000}.
4. Variation on “group.prob” for “HALLMARK COMPLEMENT”: {(0.9, 0.1), (0.8, 0.2), (0.7, 0.3), (0.6, 0.4), (0.5, 0.5), (0.4, 0.6), (0.3, 0.7), (0.2, 0.8), (0.1, 0.9)}.

### B.2 Description and Pre-processing of Real-world Datasets

Glioblastoma^3^ [56] is a scRNA-seq dataset that contains the transcriptomic data of cells from a chromosomal unstable glioblastoma cancer stem cell line (cancer). It also includes cells from a diploid neural stem cell line (neural) as normal controls. Since this glioblastoma dataset contains only 59 cancer cells and 75 neural ones, it is used to evaluate how DeepGSEA performs on scRNA-seq datasets with a small sample size.

Influenza^4^ [42] is a scRNA-seq dataset with gene profiles of A549 cells undergoing either a mock infection (4505 cells) or infection by the influenza strain PR8-NS1-GFP at MOIs of 0.2 (7377 cells) and 2.0 (6758 cells). We use this dataset to evaluate our model’s performance when dealing with multiple cell conditions. Downsampling of the dataset is performed to accelerate the GSE analysis and to evaluate the performance of DeepGSEA on a subset of cells. Specifically, 500 cells are sampled from each group.

The alzheimer^5^ [55] dataset has scRNA-seq profiles of 8186 cells collected from the brain tissues of mice with Alzheimer’s disease, with another 8506 cells from the control group. Cell type information is provided in this dataset so it can be used to test how DeepGSEA manages to leverage the additional knowledge of cell types to better examine and interpret the enrichment of gene sets.

Since the real-world datasets in our experiments have been processed differently prior to publication, we perform different preprocessing for different datasets. For the glioblastoma dataset, as it was already log-transformed, we only scale it to unit variance and zero mean and truncate values greater than 10. The cell counts in the influenza dataset have been normalized, and we take log-transformation and scaling as the preprocessing of it. We do not perform any additional preprocessing for the alzheimer dataset, because the provided dataset has already been standardized.

### B.3 Implementation Details of DeepGSEA model

In our experiments, DeepGSEA models are built with a three-layer shared encoder, and the gene set heads are fully connected layers as mentioned in Section 3.2. The hidden dimension h dim is set as 64 and the dimension of latent embeddings z dim is set as 32.

## C More Experimental Results

### C.1 DeepGSEA’s Classification on Real-world Datasets

As Formula 5 shows the final prediction is given based on a simple linear combination of information from different gene sets, the performance of cell classification can reflect how well the model captures phenotype knowledge from gene sets. Table 3 presents DeepGSEA’s performance of phenotype classification of cells in the test sets (Formula 5) on different datasets with various gene set databases. We can observe that DeepGSEA does effectively capture the knowledge of phenotypes from gene sets and distinguish different cell groups, as evidenced by an auROC score over 0.89 on all datasets. For the glioblastoma dataset, both GO and Pathway are informative enough to explain variations in gene expression of cells with different phenotypes. However, models with GO or Pathway behave differently when analyzing phenotypes in influenza and alzheimer datasets. While Pathway is better suited to the influenza dataset than GO, DeepGSEA with GO performs better than the one with Pathway on the alzheimer dataset. Generally, DeepGSEA’s performance in classifying a dataset using a given gene set database can reflect the appropriateness of that database for the analysis.

**Table 3:** DeepGSEA’s phenotype classification performance on real-world datasets (average on 5 folds). The metrics for classification reflect how well the gene sets in the database can be used to explain phenotypes of interest.

|  | glioblastoma<br>(GO) | glioblastoma<br>(Pathway) | influenza<br>(GO) | influenza<br>(Pathway) | alzheimer<br>(GO) | alzheimer<br>(Pathway) |
| --- | --- | --- | --- | --- | --- | --- |
| Accuracy | 1.00 | 1.00 | 0.78 | 0.95 | 0.80 | 0.75 |
| auROC | 1.00 | 1.00 | 0.92 | 0.99 | 0.89 | 0.84 |
| F1-score | 1.00 | 1.00 | 0.78 | 0.95 | 0.80 | 0.76 |

### C.2 Ablation Studies

We first perform ablation studies on the alzheimer dataset as it is the most challenging one in our study and can reflect how the results will change with variations to the design. As we mentioned in Section 4.3, DeepGSEA is trained with cell type labels given by the alzheimer dataset. In order to evaluate the performance of DeepGSEA on the dataset without information of cell types, we trained another model using Formula 17 instead of Formula 18 in model tuning, which we term as DeepGSEA_no_label_. In addition, DeepGSEA_one_proto_ is trained as a model variant that learns only one prototype for the distribution of each phenotype. A model variant called DeepGSEA_one_stage_ is trained to test the performance of the model without the two-stage training paradigm mentioned in Section 3.3. DeepGSEA_one_gene_set_ is a model trained only with the gene set “generation of neuron”, which is used to test if the model performance benefits from the proposed backbone-head architecture. All model variants except DeepGSEA_one_gene_set_ are trained with GO as the source of gene sets.

Table 4 shows the classification performance of the model variants, where we examine each model’s ability to classify cells based on either the GO term “generation of neurons” or all gene sets. Without the additional biological knowledge of cell types, DeepGSEA_no_label_ and DeepGSEA_one_proto_ have comparable performance to the full version of DeepGSEA in terms of phenotype classification. However, Fig. 10 (a) and (b) show that DeepGSEA_no_label_ tends to group cells of the same phenotype together in the latent space but remove their characteristic of cell types. Such a phenomenon is also presented in Fig. 10 (c) and (d), where the distributions of different phenotypes are further squashed in the latent space as there is only one prototype for each phenotype.

**Table 4:** Classification performance (auROC) of different model variants on the alzheimer dataset.

| | DeepGSEA | $\text{DeepGSEA}_{\text{no\_label}}$ | $\text{DeepGSEA}_{\text{one\_proto}}$ | $\text{DeepGSEA}_{\text{one\_stage}}$ | $\text{DeepGSEA}_{\text{one\_gene\_set}}$ |
| --- | --- | --- | --- | --- | --- |
| “generation of neurons” | 0.83 | 0.83 | 0.83 | 0.82 | 0.75 |
| all gene sets involved | 0.89 | 0.87 | 0.88 | 0.86 | 0.75 |

**Fig. 10:**
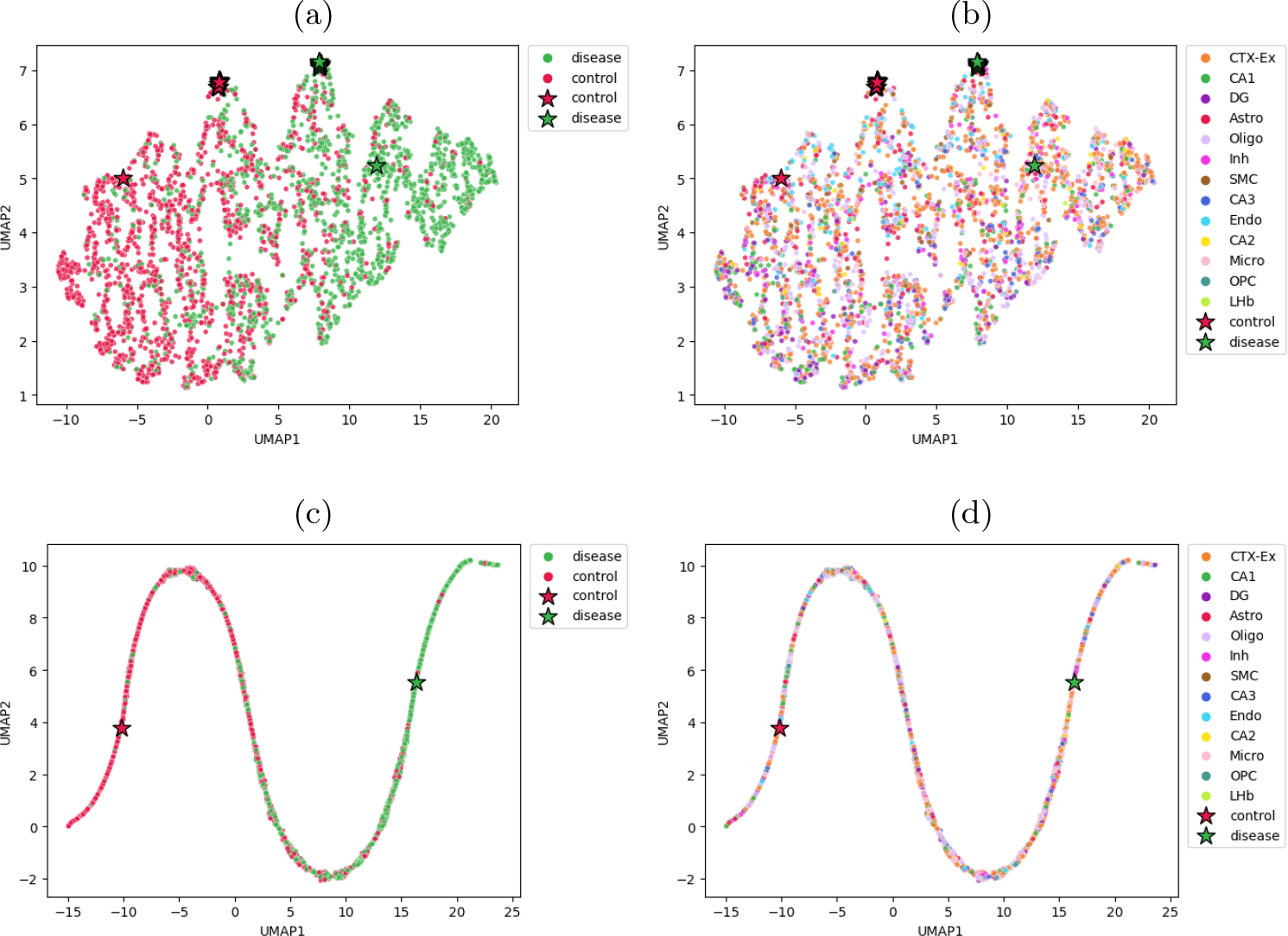
Visualized latent spaces learned by different model variants with color annotations: (a) phenotype distribution in DeepGSEA_no_label_, (b) cell type distribution in DeepGSEA_no_label_, (c) phenotype distribution in DeepGSEA_one_proto_, (d) cell type distribution in DeepGSEA_one_proto_.

In contrast, the cell types are more well reflected in the visualized latent spaces of DeepGSEA_one_stage_ and DeepGSEA_one_gene_set_ (Fig. 11 (b) and (d)), since cell type annotations are utilized in their training. But according to Table 4, DeepGSEA_one_stage_ underperforms the origin model in terms of using all gene sets for prediction as the weights to combine gene sets may be incorrectly learned when the gene set encoders are not well trained. DeepGSEA_one_gene_set_ performs even worse than all other model variants on the classification in both cases, suggesting that the backbone-head architecture in DeepGSEA does help the model to better encode each gene set by sharing the knowledge about other gene sets.

**Fig. 11:**
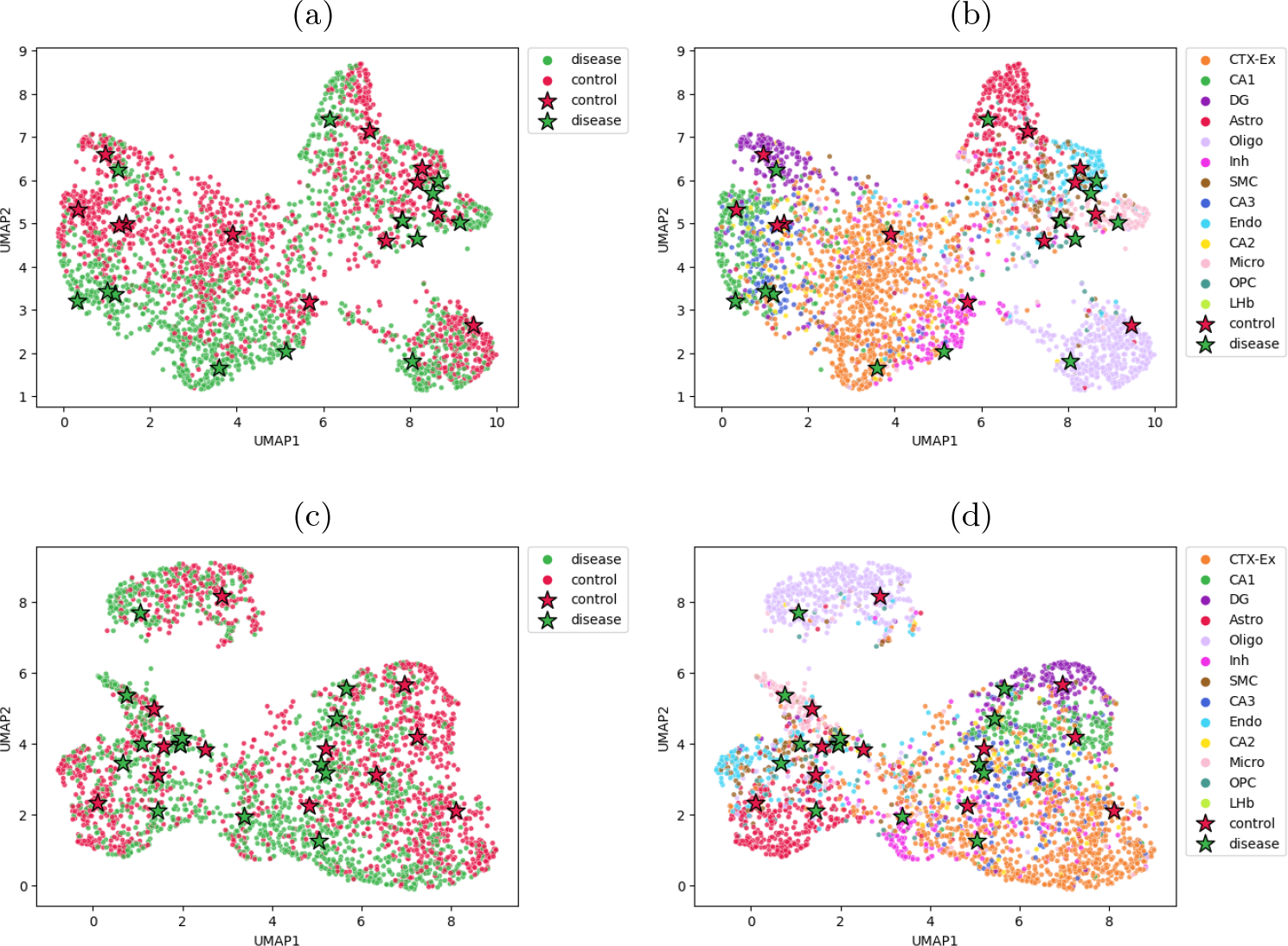
Visualized latent spaces learned by different model variants with color annotations: (a) phenotype distribution in DeepGSEA_one− stage_, (b) cell type distribution in DeepGSEA_one− stage_, (c) phenotype distribution in DeepGSEA_one− gene− set_, (d) cell type distribution in DeepGSEA_one− gene_ −_set_.

We also perform ablation studies on the selection of hyperparameters {*λ*_3_, *λ*_4_, *λ*_5_}, which are used to regularize the latent space and improve the quality of learned prototypes. The influenza dataset is chosen, as it is simpler than the alzheimer dataset so that we can clearly identify differences in the learned prototypes when perturbing the hyperparameters. We set the number of prototypes for each phenotype as 2 to examine the effect of different *λ*_3_ on the learned latent space. Our default setting for these hyparameters is *λ*_3_ = *λ*_4_ = *λ*_5_ = 1.

Fig. 12 shows how the distances between prototypes change according to the selection of *λ*_3_. A *λ*_3_ with either too small or too large value will result in unwanted outcomes. The results for variations on *λ*_4_ are presented in Fig. 13, which shows the cells are moving forward to the corresponding prototypes when *λ*_4_ gets larger, though the estimated Gaussian distributions may be quite unnatural if *λ*_4_ is too large. Fig. 14 shows the change of learned latent spaces with respect to different *λ*_5_. As the loss related to *λ*_5_ guides the movement of prototypes, by comparing different visualized latent spaces in Fig. 14, we can observe that prototypes of the “Mock” group are becoming more distant from the ones of the “MOI0.2” group when *λ*_5_ becomes larger. In addition, by comparing *λ*_5_ = 0 with *λ*_5_ ≠ 0, we notice that the prototypes tend to move to centers of regions with a higher cell density. However, if *λ*_5_ is too large, the model may fail to capture subtle differences between cells at the boundaries of two subpopulations, as the prototypes are largely determined by the center of the whole group, which can result in a missing of details at the boundaries.

**Fig. 12:**
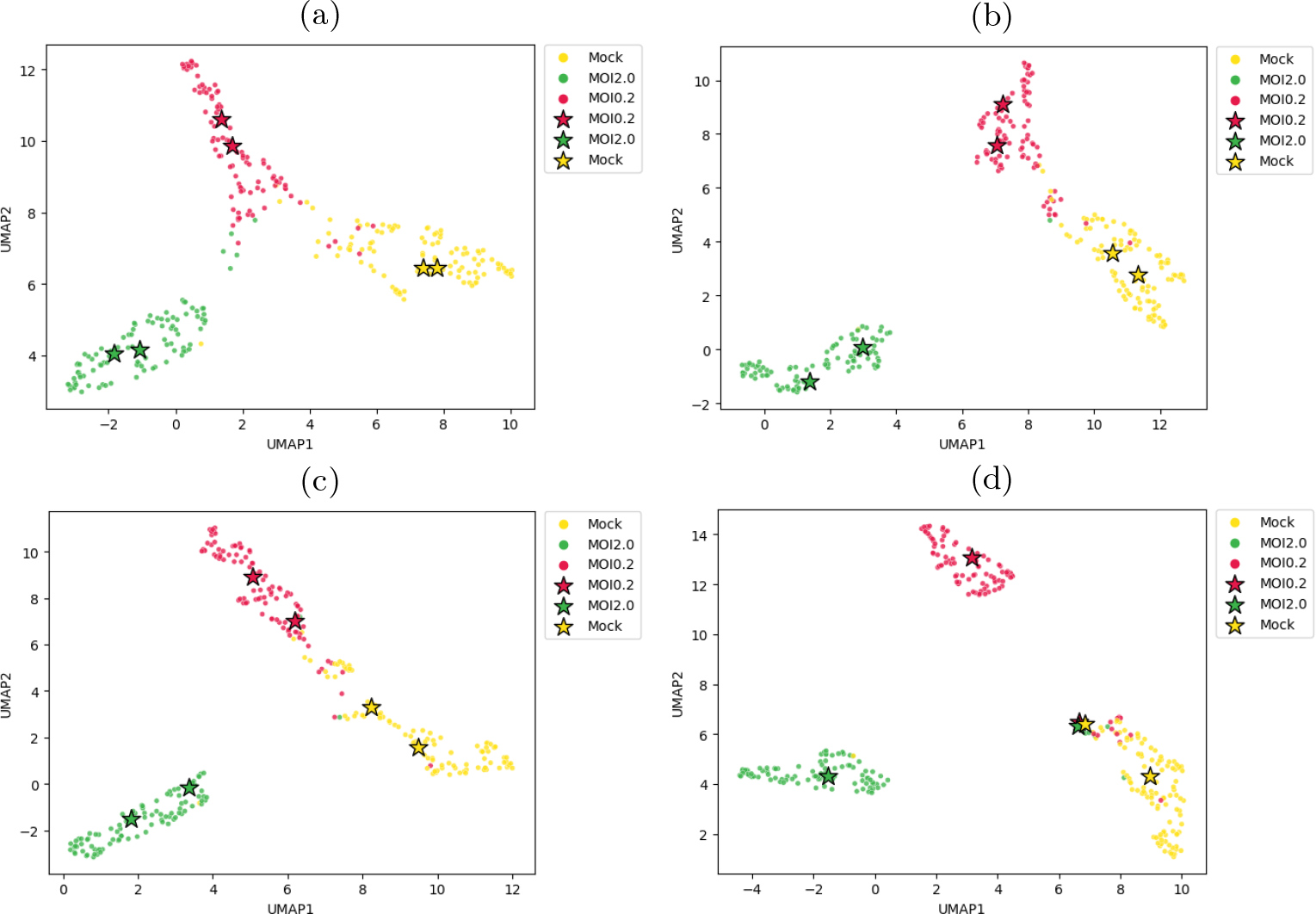
Visualized latent space of “translation” in the influenza dataset with variations on the hyperparameter *λ*_3_: (a) *λ*_3_ = 0, (b) *λ*_3_ = 1, (c) *λ*_3_ = 10, (d) *λ*_3_ = 100.

**Fig. 13:**
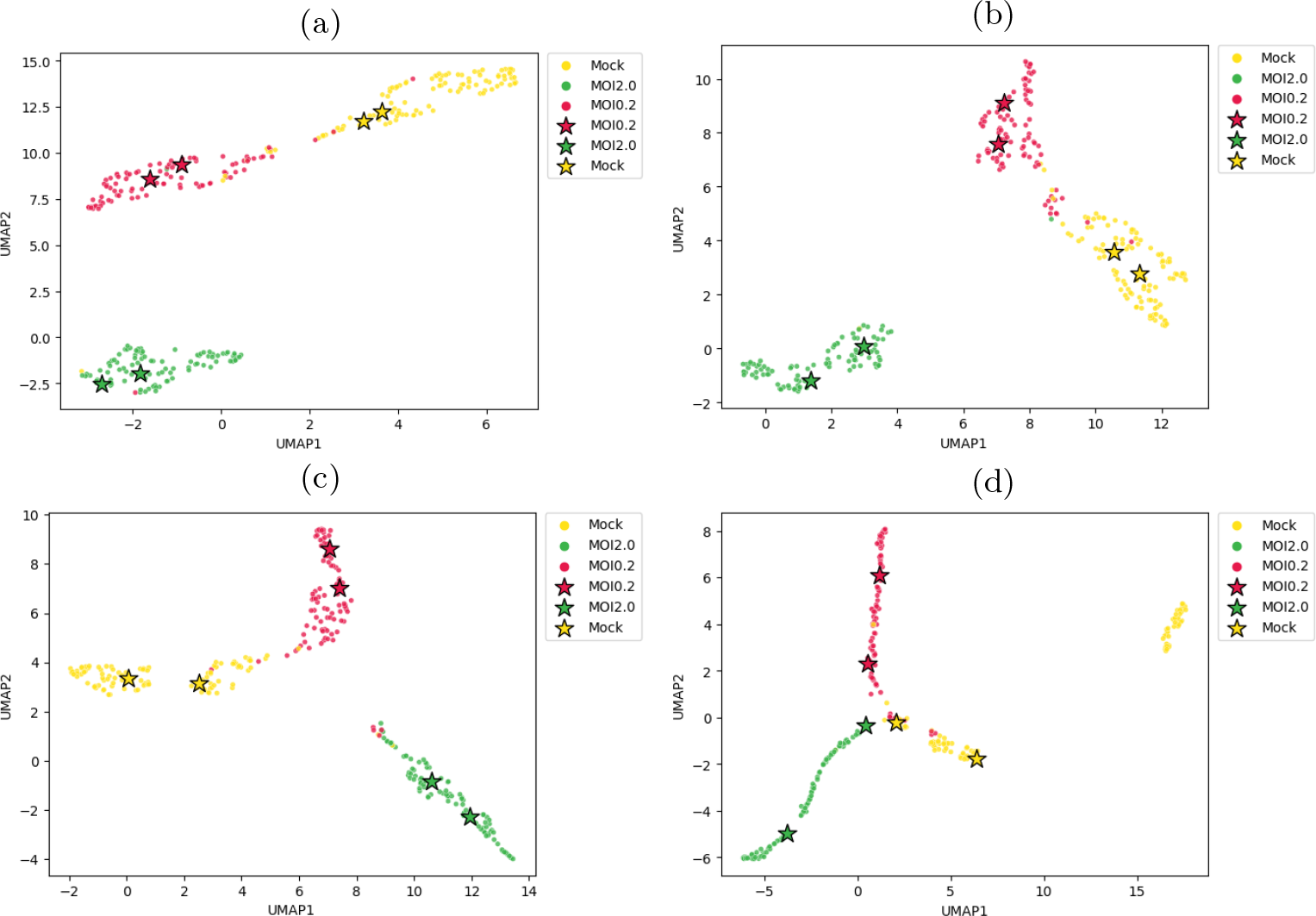
Visualized latent space of “translation” in the influenza dataset with variations on the hyperparameter *λ*_4_: (a) *λ*_4_ = 0, (b) *λ*_4_ = 1, (c) *λ*_4_ = 10, (d) *λ*_4_ = 100.

**Fig. 14:**
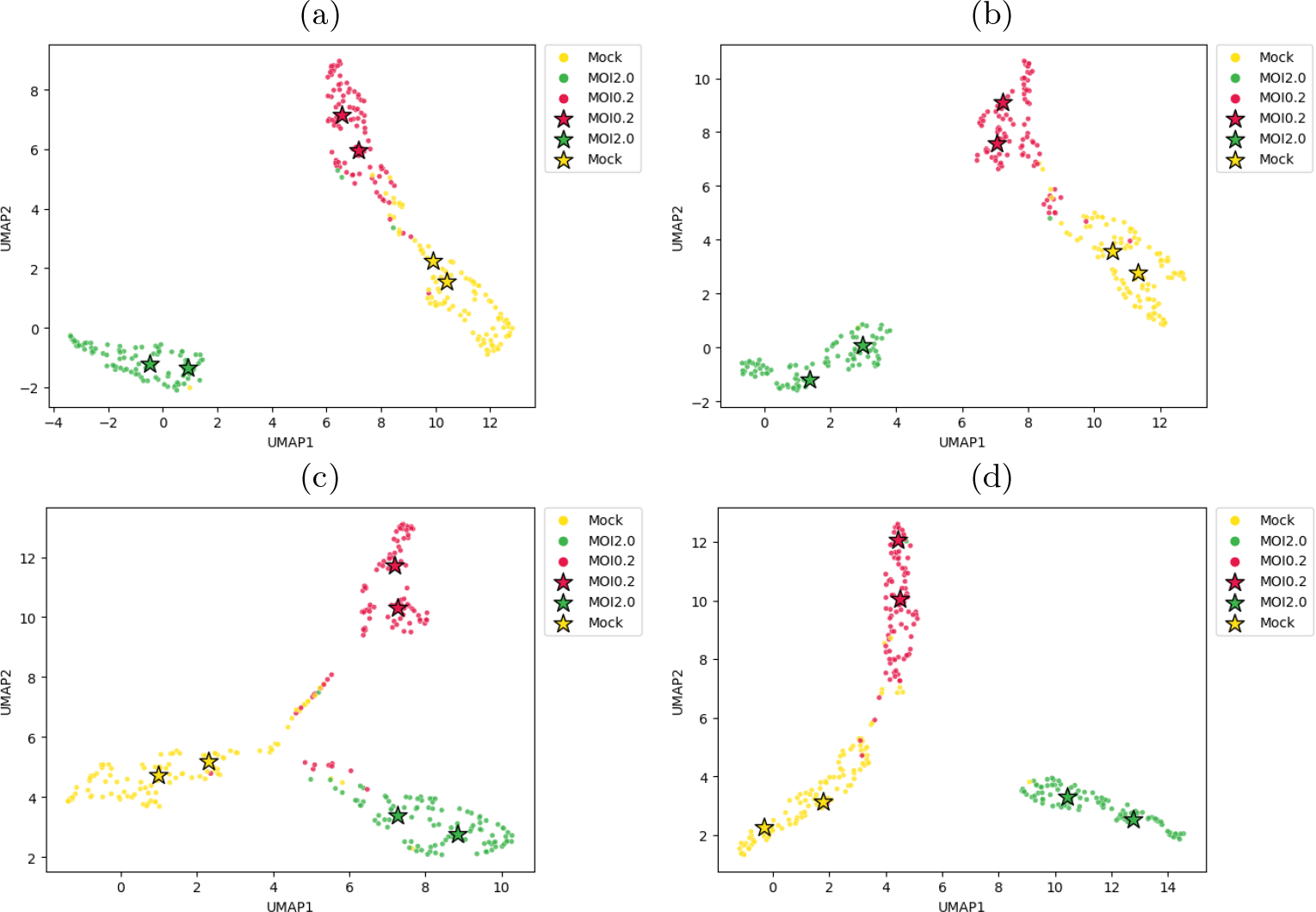
Visualized latent space of “translation” in the influenza dataset with variations on the hyperparameter *λ*_5_: (a) *λ*_5_ = 0, (b) *λ*_5_ = 1, (c) *λ*_5_ = 10, (d) *λ*_5_ = 100.

## Footnotes

1 The original data is available at https://www.ncbi.nlm.nih.gov/geo/query/acc.cgi?acc=GSE212270.

2 The definition of all parameters can be found at https://www.bioconductor.org/packages/devel/bioc/vignettes/splatter/inst/doc/splat_params.html.

3 The glioblastoma dataset is available at https://www.ncbi.nlm.nih.gov/geo/query/acc.cgi?acc=GSE132172.

4 The influenza dataset is available at https://www.ncbi.nlm.nih.gov/geo/query/acc.cgi?acc=GSE122031.

5 The alzheimer dataset is available at https://singlecell.broadinstitute.org/single_cell/study/SCP1375.

